# Identifying functionally distinctive and threatened species

**DOI:** 10.1101/2022.11.28.518165

**Authors:** Sandrine Pavoine, Carlo Ricotta

## Abstract

Functional traits determine species’ responses to environmental change and/or determine species’ effects on ecosystem functions. When species with distinctive functional traits are threatened, there is a risk that ecosystem properties are also threatened. This is because functionally distinctive species may be those that have irreplaceable roles in an ecosystem and/or those that would be able to survive unusual environmental disturbances. To include functional distinctiveness as a criterion in conservation strategies, we need formal quantification of the degree of distinctiveness and threat a species exhibits. Starting with previous quantification attempts, we develop a framework that links different viewpoints on functional distinctiveness and accounts for all species’ extinction probabilities. Our framework is particularly relevant at the local scale where species extinctions impact ecosystem functioning and where conservation policies are developed. As a case study, we thus applied our framework to the mammals of Indian dry forests known to be threatened with a drastic decrease in functional diversity. We notably highlight that although some of the functionally distinctive and threatened species we identified, such as the tiger, are charismatic and considered by conservation actions, others are not. This is the case for some rat species and pangolins, whose negative image in the media during the COVID-19 pandemic could be detrimental to attracting public interest in their preservation. From this case study, we note that noncharismatic, less known species that may be key for ecosystems could be revealed by applying our framework to a range of ecosystems and taxa.

## 1. Introduction

Traits can reflect a species’ ecological strategies and its function in its ecosystem (Cooke et al., 2020). Traits considered functional may indeed represent species’ morphological, physiological, phenological or behavioral features (e.g., Carmona et al., 2021) that determine the effect a species has on its ecosystem properties (“effect traits”, e.g., Lavorel and Garnier, 2002; Gorczynski et al., 2021). This definition is mainly shared in studies of functional distinctiveness, where the main objective is to evaluate the ecological impacts species extinction may have in the near future on ecosystems (e.g., Fritz and Purvis, 2010; Toussaint et al., 2021). The other facet of functional traits is the species’ sensitivity or response to environmental changes (“response traits”). Species that have distinctive response-trait values and redundant effect traits may ensure the stability of ecosystem functioning if their distinctive response traits allow them to cope better than other species with some specific environmental change (response-trait diversity as insurance, Loreau et al., 2001). Redundancy in response trait values within a species assemblage would, in contrast, lead to high extinction risk and a drastic decline in effect trait diversity in the case of environmental disturbance. Functionally distinctive species thus globally ensure the maintenance of “nature’s variability”, contributing to ecological diversity, which is indispensable for ecosystem processes and functions (Cooke et al., 2020).

The loss of functionally distinctive species may thus increase the risk that a system collapses compared to random extinction (e.g., Flynn et al., 2009). Measuring to what degree a species is functionally distinctive allows the identification of species with irreplaceable ecological strategies (Cooke et al., 2020). Then, determining those functionally distinctive species that are threatened with extinction would allow alerting conservation actors on the need to consider these species in their programs. Functional distinctiveness can be measured using a single functional trait. For example, with a quantitative trait, Redding et al. (2010) used the absolute distance from the average or median score for a trait as an index of distinctiveness. With a categorical trait, they used the frequency of species that share the same category as the target species as an index of ordinariness (i.e., this frequency decreases with the target species’ distinctiveness). When several traits are considered, however, those simple measurements are not used. In that case, the calculation of functional dissimilarities between species has the advantage of allowing the consideration of any type and any number of traits when comparing species (e.g., Pavoine et al., 2009).

Once functional dissimilarities have been calculated, the most commonly used indices of functional distinctiveness are the average functional dissimilarity to other species and the minimum functional dissimilarity to another species. These two indices may, however, lead to distinct priority settings (e.g., Cooke et al., 2020). Indeed, a species may be functionally distant, on average, from all other species but functionally similar to one of them (Kosman et al., 2019). The mathematical index used (e.g., average versus minimum dissimilarity) is thus likely to influence the conclusions on which species are most functionally distinctive and thus the decisions in terms of conservation priority. Pavoine and Ricotta (2021) thus developed an approach that identifies the definition(s) of functional distinctiveness (from the distance to the most functionally similar species, through classical average dissimilarity to other species, to the maximum dissimilarity with another species) according to which the conservation priority for a species could be decided. Here, we extend this approach to allow for the consideration of species extinction probabilities in functional distinctiveness. This extended framework allows the identification of those species that are of conservation interest according to a large range of viewpoints on functional distinctiveness.

As an illustration, we analyzed the functional distinctiveness of mammals in the dry forests of India. Most analyses of functional distinctiveness have been performed globally (at the world scale), while local analyses may be the most impactful for three reasons (Toussaint et al., 2021). First, most conservation policies are defined at a local, country level within nations (see also, e.g., Brodie et al., 2021). Second, as countries host distinct types of ecosystems and faunas, small scales are those where the impacts of extinction on ecosystem functioning can be studied (see also, e.g., Carmona et al., 2021). Third, a species may be functionally distinctive in a country, possibly fulfilling a critical role in its ecosystem, while its trait values could be more frequent elsewhere so that it would not have been considered functionally distinctive at a global scale. We chose to focus on the Indo-Malay realm because it exhibits high functional diversity in vertebrates and is susceptible to drastic loss in functional diversity in the near future (Fritz and Purvis, 2010; Toussaint et al., 2021); the most drastic loss of all realms if currently threatened mammal species were driven extinct (Toussaint et al., 2021). Within the Indo-Malay realm, India and the forest biome exhibit the highest number of threatened species (Redford et al., 2011). In this context, the mammals of the Indian dry forests, with high species richness and functional diversity, appeared to us as a relevant group to analyze for our illustration.

## 2. Methods

We start here with an index of functional distinctiveness named 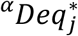 (Pavoine and Ricotta, 2021) that relies on functional dissimilarities among species. Then, we add to this index information on species extinction risk.

Consider an assemblage of *N* species. Let *d*_*ij*_ be the functional dissimilarity between *i* and *j*. Consider that the *N* species are ordered from the most similar to the most dissimilar species to *j*. Of course, the species that is most similar to *j* is *j* itself (*d*_*jj*_ = 0); we denote *c*, the *c*^*th*^ species most functionally similar to *j* and pose *c* = 0 for species *j* itself. *c* thus varies from 0 to *N*-1. Note that the framework is still valid if there are ties in the *d*_*ij*_ values and the species that have even functional dissimilarities to *j* can be ordered in any way without changing the value of 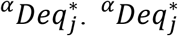 is a parametric index of functional distinctiveness, where the parameter *α* varies from -∞ to +∞. Compared to the Pavoine and Ricotta (2021) formulation of 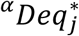, we reformulate 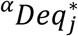 as follows (see Appendix A for the original formula):

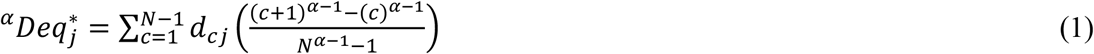

Considering the limit of 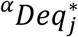 when *α* tends to 1, we pose 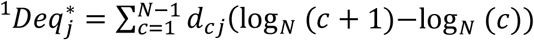.

Our new formulation of index 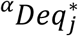 explicitly highlights that parameter *α* controls the importance given to species with close trait values compared to those with distant trait values in the calculation of functional distinctiveness. With Eq. 1, 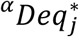 is revealed to be the weighted mean of the functional dissimilarities between species *j* and all other species with weights equal to

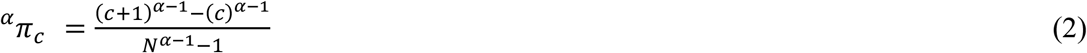

knowing that 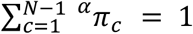 (Appendix A). The case *α* = 2 is the only case for which species’ weights are even (^*α*^ *π*_*c*_= 1/(*N*-1)). When *α* < 2, species weights increase from the species most functionally dissimilar to the species least functionally dissimilar to *j*. The lowest *α* is, the steepest the increase in weights, so that when *α* tends to -∞, 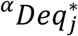 tends to min_*i*_(*d*_*ij*_). Conversely, with *α* > 2, species weights increase from the species least functionally dissimilar to the species most functionally dissimilar to *j*. The highest *α* is, the steepest the increase in weights, so that when *α* tends to +∞, 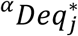 tends to max_*i*_(*d*_*ij*_) (Appendix B in Pavoine and Ricotta, 2021) (see Fig. 1 for a theoretical example). Our reformulation of 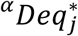 thus allows an easier interpretation of the values taken by this index.

**Fig. 1.**
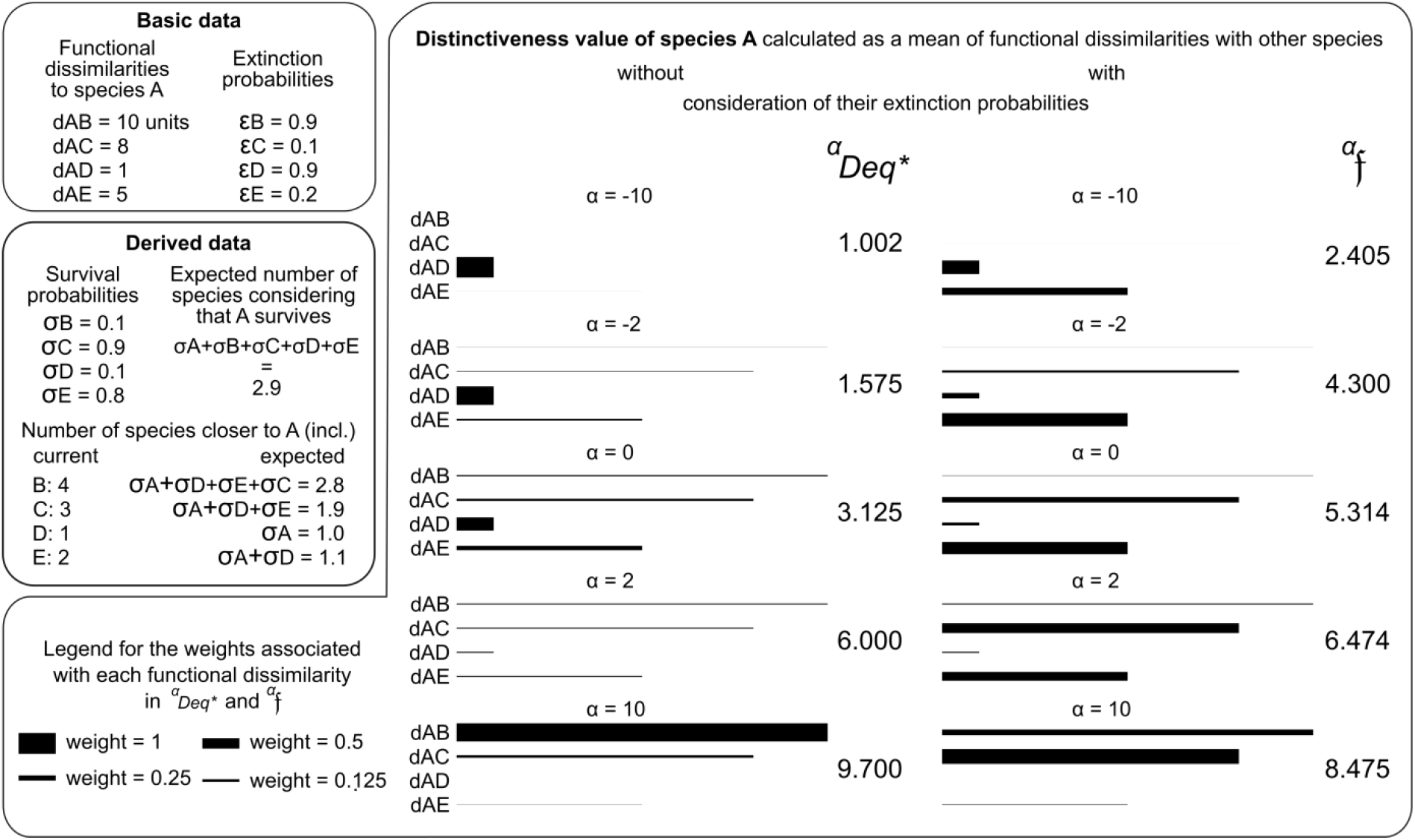
Theoretical example of the calculation of functional distinctiveness as a mean of functional dissimilarities with other species. Basic data specify the functional dissimilarities (d) and extinction probabilities (ε). The derived data specify, for each species, its survival probability (σ = 1-ε) and its order of proximity with species A (number of current species closer to A). The derived data also provide the total expected number of species and the expected number of species that are functionally closer to A (including A) than the specified species, both calculated considering that species A survives (σ_A_ = 1). In the largest box on the calculation of the distinctiveness of A, the length of a segment represents a value of functional dissimilarity as specified in the basic data. The thickness of a segment represents the weight used in ^*α*^*Deq\** and ^*α*^*f*, both being weighted means of functional dissimilarities. The absence of a segment means that the weight is negligible (<<0.001) so that the corresponding functional dissimilarity hardly influences the distinctiveness value. For each index (^*α*^*Deq\** and ^*α*^*f*), distinctiveness values are given for 5 values of their parameter *α*, and they are rounded to 3 decimal places.

The main novelty of our work is the inclusion of species extinction probabilities in Eq. 1. Let *ε*_*i*_ and *σ*_*i*_ = 1 − *ε*_*i*_ be the extinction probability and survival probability of any species *i*, respectively. As in Pavoine and Ricotta’s (2022) framework on evolutionary distinctiveness, here for functional distinctiveness we consider that the target species *j* is safe, and we replace *N* with *N*_*j*_, the sum of the survival probabilities of all species (i.e., 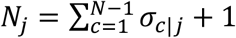), and *c* with *N*_*c-1*|*j*_, the sum of the survival probability of species *j* and of the *c-1* other species that are the most functionally similar to species *j* (i.e., for *c* = 1, *N*_*c*-1|*j*_ = 1, survival probability of species *j*, and for 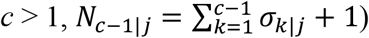) (Fig. 1). This yields:

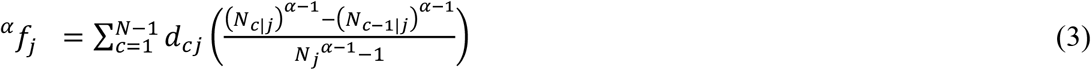

^*α*^*f*_*j*_ is a weighted mean. In Eq. 3, the functional dissimilarity between *j* and the *c*^*th*^ species most functionally similar to *j* is indeed weighted by

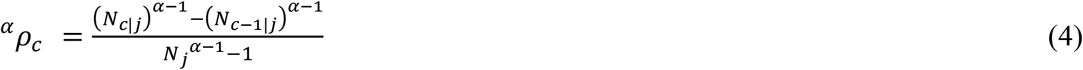

with 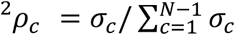.

To complement ^*α*^*f*_*j*_, we define 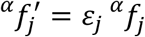 and 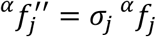 so that 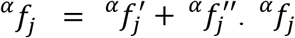 allows the identification of functionally distinctive species, 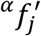 allows the identification of threatened functionally distinctive species, and 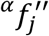 allows the identification of safe functionally distinctive species.

## 3. Case study

### 3.1. Data

We considered 94 mammal species that occur in Indian dry forests. This amounts to all species listed in the International Union for the Conservation of Nature Red List (IUCN, 2022) in the dry forests of India except one of them, *Belomys pearsonii*, whose extinction risk is unknown. We considered six traits: body mass (g), diet (percentage of consumption across plant, invertebrate and vertebrate food items), foraging stratum (with three possibilities: ground level including aquatic foraging, arboreal and scansorial), activity cycle (nominal trait with three levels: 1=nocturnal only, 2=crepuscular or cathemeral, 3=diurnal only), habitat breadth (number of distinct suitable level 1 IUCN habitats from 1 to 7), use of freshwater habitat (with two possibilities, yes or no) (data obtained from Soria et al., 2021, except activity cycle for *Nesokia indica* and *Tetracerus quadricornis* obtained from Sridhara and Tripathi, 2005 and Meghwal et al., 2018). Adult body mass data were cube-root transformed before analysis. This transformation led to a linear relationship between body mass and body size (Appendix B; Fig. B.1.) and avoided the spurious assumption that the biological impact of a given difference in body mass would be the same between two small species as between two large species (Fritz and Purvis, 2010). We did not use body size, as some values were missing for this trait. Contrary to Fritz and Purvis (2010) and Cooke et al. (2020), we did not log-transform body mass, as this would have led to a bell-shaped distribution with a drastic undervaluation for the distinctiveness of large mammals. We obtained extinction probabilities within the next 100 years with the model of Andermann et al. (2021; Crit E EX mode) that considers each species’ current conservation status (IUCN, 2022), chances of conservation status changes (with the mammals as a reference group), and generation length (obtained from Pacifici et al., 2013; completed by data available in Soria et al. (2021) for *Catopuma temminckii* and *Herpestes auropunctatus*).

### 3.2. Analysis

To ease the illustration of our framework, we first focused on a single functional trait. We chose cube-root transformed body mass, as body mass, or the related body size, is systematically included in analyses of functional distinctiveness (e.g., Leitão, 2016; Cooke et al., 2020; Toussaint et al., 2021). Indeed, “size or body mass is an important predictor of many ecological traits in mammals and an indicator of a species’ ecological niche” (Fritz and Purvis, 2010). It has a broad range across mammals and underlines many physiological and ecological processes (e.g., predation and seed dispersal; Cooke et al., 2020).

We used Pavoine et al.’s (2009) mixed-variables coefficient of distance to calculate functional distances from cube-root transformed body mass and then all functional traits listed above (underlying metrics were the Manhattan metric for quantitative and ordinal traits, the Gower (1971) metric for nominal traits, and the Orloci (1967) chord distance for the diet expressed in terms of percentage of use). We then identified species that were in the top-^*α*^*Deq*\*, top-^*α*^*f*, top-^*α*^*f*′, and top-^*α*^*f* ′′, which we defined as the set of species that were among the 10th most distinctive according to ^*α*^*Deq*\*, ^*α*^*f*, ^*α*^*f*′, and ^*α*^*f* ′′, respectively, for at least one value of *α* we considered. For these calculations, we considered *α* varying from - 10 to 10 with a step of 0.1.

## 4. Results

### 4.1. Body mass distinctiveness

Considering body mass only, the top ^*α*^*Deq*\* was composed of the 23 species listed in Fig. 2. Among them, the two species with the highest body mass were classified first (elephant, *Elephas maximus*) and second (rhino, *Rhinoceros unicornis*) most distinctive over all values of *α* we considered. Some were top distinctive for low values of *α* (e.g., the gazelle *Gazella bennettii*, the wild dog *Cuon alpinus*, and the wolf *Canis rufus*, even if classified among the 10th least distinctive for high values of *α*; Fig. 2). Others were top distinctive only for high values of *α* (e.g., the shrew, *Crocidura vorax* and the mouse *Vandeleuria oleracea*, even if classified among the 10th least distinctive for *α* close to zero; Fig. 2). Some were top distinctive for *α* of approximately 2 but not for high or low values of *α* (e.g., the wild boar *Melursus ursinus*).

**Fig. 2.**
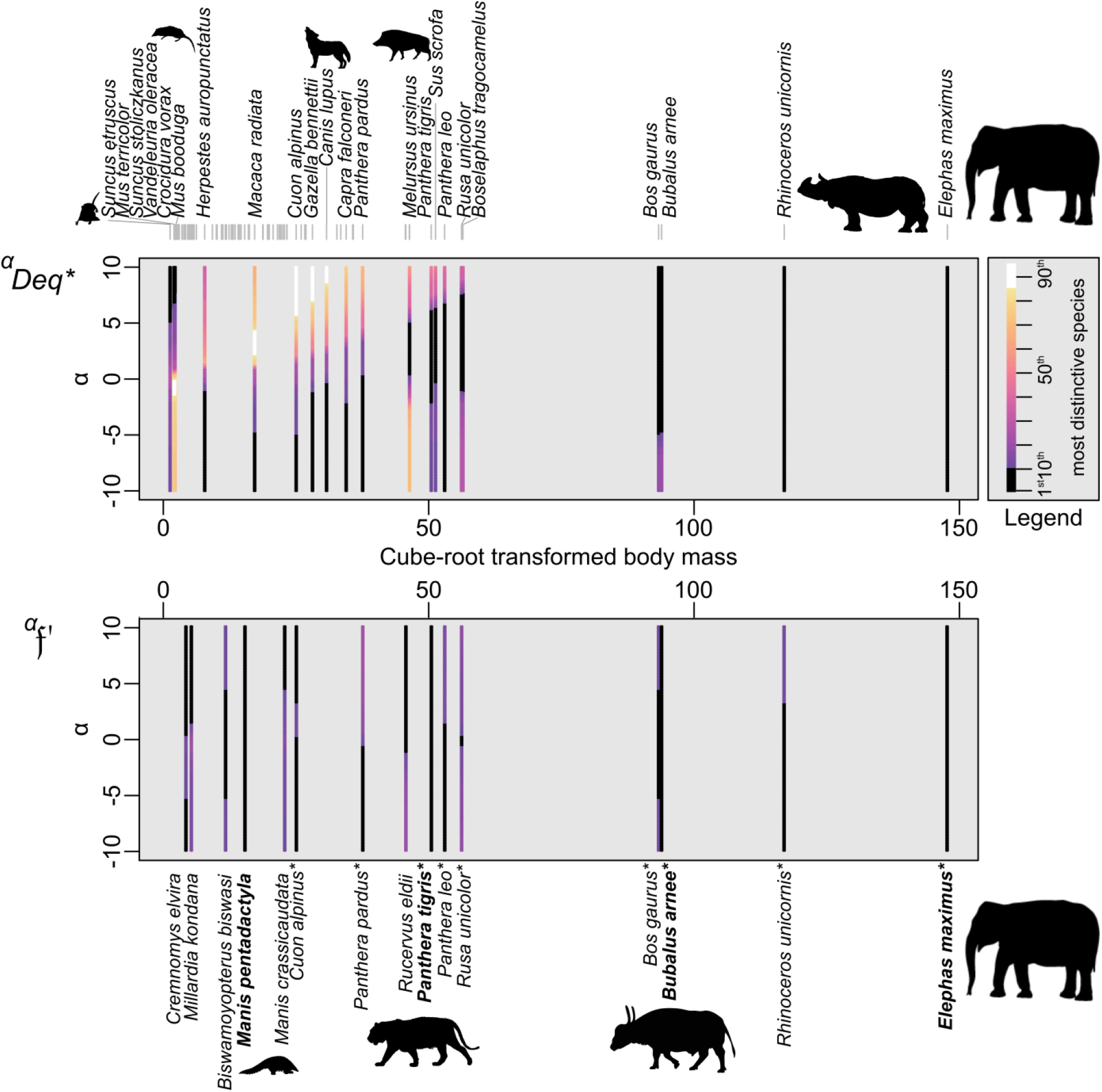
Distinctiveness variation across *α* values for top-^*α*^*Deq\** species and top-^*α*^*f*′ species (i.e., species that are at least once among the 10 most distinctive according to ^*α*^*Deq\** and ^*α*^*f*′, respectively). The names of species in the top-^*α*^*Deq\** are indicated on the top of the figure, and those of species in the top-^*α*^*f*′ are indicated on the bottom of the figure. Species names in bold indicate the four species that have been classified among the 10 most distinctive species according to ^*α*^*f*′ for all values of *α* we considered. The x-axis provides the species’ body mass, and the y-axis provides the value of parameter *α* in ^*α*^*Deq\** and ^*α*^*f*′ Colors specify the distinctiveness rank from black (the 10 most distinctive species) to white (the 10 least distinctive species). Gray segments at the top of the figure indicate body mass for all 94 species.

Adding extinction probabilities, we observed that species’ ranks changed (except for *E. maximus* and *R. unicornis*) but not enough to affect the list of top distinctive species except for one species identified in the top-^*α*^*f* but not in the top-^*α*^*Deq*\* (Appendix B, Fig. B.2): the mainland serow (*Capricornis sumatraensis*), a Bovidae species that has a small part of its distribution range in the northern forest of India on the slopes of Himalaya. *C. sumatraensis* is the species with the greatest rank change between ^*α*^*Deq*\* and ^*α*^*f* (Appendix B, Fig. B.2).

Accounting for a species’ own extinction probability, only nine species out of the 24 species detected as distinctive with ^*α*^*Deq*\* and ^*α*^*f* were also among the top-^*α*^*f*′ distinctive and threatened species: *Bos gaurus, Bubalus arnee, C. alpinus, E. maximus, Panthera leo, Panthera pardus, Panthera tigris, R. unicornis*, and *Rusa unicolor* (Fig. 2). The top ^*α*^*f*′ list was also composed of 6 additional species: *Biswamoyopterus biswasi, Cremnomys elvira, Manis crassicaudata, Manis pentadactyla, Millardia kondana*, and *Rucervus eldii*. Only four species were classified among the 10 most distinctive and threatened across all values of *α* (Fig. 2): the Asian elephant *E. maximus* (from 1st to 6th highest priority depending on *α*), the wild water buffalo *B. arnee* (from 1st to 10th), the Chinese pangolin *M. pentadactyla* (always 8th), and the tiger *P. tigris* (from 5th to 10th).

Accounting for species’ own survival, instead of extinction probability, index ^*α*^*f* ′′ revealed that all species in top-^*α*^*f* ′′ belong to the list of top-^*α*^*f* and top-^*α*^*Deq*\* (thus having the highest distinctiveness values, independent of their own extinction risk). There was one exception: *Ursus thibetanus*, with a relatively high survival probability, σ = 0.88, classified as having the 10th highest value for ^*α*^*f* ′′ (with α in [0.2, 4.1]), while it had the 11th highest value for ^*α*^*f* (with α in [0.2, 4.1]) and ^*α*^*Deq*\* (with α in [0.4, 5]). Additionally, analyzing the index ^*α*^*f* ′′ revealed that the distinctiveness in body mass was so high for some species that these were both in the top ^*α*^*f* ′′ and in the top ^*α*^*f*′ : *B. gaurus, B. arnee, E. maximus, P. leo, P. pardus, P. tigris, R. unicornis*, and *R. unicolor*. However, they were not classified among the 10 highest values for the same values of parameter α.

### 4.2. Multitrait distinctiveness

Adding information on other traits, the lists of top-^*α*^*Deq*\* and top-^*α*^*f* species were identical to each other but different from those obtained by considering body mass only (Table 1). Only the four largest species (*E. maximus, R. unicornis, B. arnee, B. gaurus*) and the leopard (*P. pardus*) appeared as top distinctive both when we considered body mass only and when we considered a larger range of traits. When accounting for all traits, *Neofelis nebulosa* (the diurnal clouded leopard, a relatively large Carnivora species with scansorial foraging and restricted habitat breadth) and *R. eldii* (the crepuscular herbivorous Eld’s deer with large habitat breadth, including wetlands, and restricted in India to Keibul Lamjao National Park, Eliza et al., 2018) (IUCN, 2022) were top-^*α*^*Deq*\* and top-^*α*^*f* across all the *α* values considered. A large part (2/3) of the top-^*α*^*Deq*\* and top-^*α*^*f* species were also in the top-^*α*^*f* ′′ list. Some with high distinctiveness (^*α*^*f*) values for some (*B. gaurus, E. maximus, P. pardus*) or all (*N. nebulosa*) *α* were both in the top-^*α*^*f*′ and top-^*α*^*f* ′′ list.

**Table 1.** List of top-^*α*^*Deq\** and top-^*α*^*f*′ species when accounting for all traits (body mass, activity cycle, diet, foraging stratum, habitat breadth and use of freshwater habitat). Here, we indicate, across the *α* values we considered (from -10 to 10 with a step of 0.1), the *α* values (named top-10 *α*) for which the species was ranked among the 10 most distinctive (for top-^*α*^*Deq\**) and the 10 most distinctive and threatened (for top-^*α*^*f*′) species, the highest and lowest priority rates that the species obtained, and whether the species was also classified in the lists of top-^*α*^*f* and top-^*α*^*f* ′′ species (a cross means that it was).

| | Top-10 $\alpha$ | Highest priority | Lowest priority | Top-$^{\alpha}f$ | Top-$^{\alpha}f''$ |
| --- | --- | --- | --- | --- | --- |
| Always top-$^{\alpha}Deq^*$ | | | | | |
| *Neofelis nebulosa* | all | 1st | 4th | × | × |
| *Rucervus eldii* | all | 3rd | 6th | × |  |
| Top-$^{\alpha}Deq^*$ for high $\alpha$ | | | | | |
| *Prionailurus viverrinus* | > -1.9 | 1st | 14th | × |  |
| *Rhinoceros unicornis*<sup>1</sup> | > 0 | 1st | 21st | × | × |
| *Lutra lutra* | > -0.9 | 3rd | 15th | × | × |
| *Bubalus arnee*<sup>1</sup> | > 0.7 | 4th | 22nd | × |  |
| *Loris lydekkerianus* | > 1.2 | 7th | 24th | × | × |
| *Hoolock hoolock* | > 2.1 | 8th | 52nd | × |  |
| Top-$^{\alpha}Deq^*$ for restricted ranges of $\alpha$ | | | | | |
| *Prionailurus rubiginosus* | [0.3, 2.6] & > 4.5 | 7th | 16th | × | × |
| *Bos gaurus*<sup>1</sup> | [-8.4, 0.7] | 7th | 27th | × | × |
| *Macaca assamensis* | [2.7, 5.6] | 9th | 84th | × | × |
| *Manis crassicaudata* | [1.9, 2.1] | 10th | 25th | × |  |
| *Macaca leonina* | [3, 4.5] & [5.7, 10] | 10th | 85th | × |  |
| Top-$^{\alpha}Deq^*$ for low $\alpha$ | | | | | |
| *Moschiola indica* | < 1.3 | 1st | 18th | × | × |
| *Martes flavigula* | < 3.0 | 2nd | 18th | × | × |
| *Manis pentadactyla* | < 1.9 | 5th | 18th | × |  |
| *Paradoxurus hermaphroditus* | < 0.3 | 5th | 49th | × | × |
| *Elephas maximus*<sup>1</sup> | < -0.8 | 6th | 33rd | × | × |
| *Panthera pardus*<sup>1</sup> | < 1.8 | 8th | 33rd | × | × |
| *Prionailurus bengalensis* | < 0.1 | 8th | 36th | × | × |
| *Mellivora capensis* | < -8.4 | 10th | 46th | × | × |
| Always top-$^{\alpha}f'$ | | | | | |
| *M. pentadactyla*<sup>2</sup> | all | 1st | 1st | × |  |
| *R. eldii*<sup>2</sup> | all | 2nd | 4th | × |  |
| *M. crassicaudata*<sup>2</sup> | all | 2nd | 6th | × |  |
| *B. arnee*<sup>2</sup> | all | 4th | 8th | × |  |
| *Panthera tigris*<sup>2</sup> | all | 6th | 9th |  |  |
| Top- $\alpha f'$ for high $\alpha$ | | | | | |
| *Biswamoyopterus biswasi*<sup>2</sup> | > -1.2 | 2nd | 16th |  |  |
| *Cremonomys elvira*<sup>2</sup> | > -0.2 | 3rd | 26th |  |  |
| *Millardia kondana*<sup>2</sup> | > 0.5 | 5th | 26th |  |  |
| *Cuon alpinus*<sup>2</sup> | > -1.1 | 7th | 15th |  |  |
| *H. hoolock* | > 0.9 | 10th | 22nd | × |  |
| Top- $\alpha f'$ for low $\alpha$ | | | | | |
| *E. maximus*<sup>2</sup> | < -0.1 | 4th | 15th | × | × |
| *Neofelis nebulosa* | < 0.8 | 5th | 12th | × | × |
| *P. viverrinus* | < 1 | 6th | 11th | × |  |
| *P. pardus*<sup>2</sup> | < -2.1 | 9th | 18th | × | × |
| *B. gaurus*<sup>2</sup> | < -1 | 9th | 19th | × | × |

<sup>1</sup> Species that were also in the body-mass top- $\alpha Deq^*$ ; <sup>2</sup> Species that were also in the body-mass top- $\alpha f'$

Enlarging the set of traits yielded the addition of three species in the top-^*α*^*f*′ list of priority species (Table 1): *Hoolock hoolock, N. nebulosa*, and *Prionailurus viverrinus*. Conversely, three species disappeared from the top-^*α*^*f*′ list: *P. leo, R. unicolor* and *R. unicornis*. We still observed that the list of top-^*α*^*f*′ species varied according to the *α* value considered, with only 5 species in the top-^*α*^*f*′ regardless of *α*: the pangolins *M. pentadactyla* and *M. crassicaudata*, Eld’s deer (*R. eldii*), the wild water buffalo (*B. arnee*) and the tiger (*P. tigris*).

## 5. Discussion

We developed a unifying framework for the measurement of functional distinctiveness and applied this framework to detect functionally distinctive and threatened mammals in the dry forests of India. Here, we discuss the definition of functional distinctiveness in terms of available, selected data and in terms of the measurement formula. We also discuss the potential and consequences that functional distinctiveness may have as a tool for conservation policies using the example of mammals in Indian dry forests as a case study.

### 5.1. Functional distinctiveness depends on the selection of traits and a model of extinction risk

The first step when studying functionally distinctive and threatened species is to select relevant traits. This selection could indeed influence conservation priorities that rely on functional distinctiveness. Among the currently available information on species traits, organism size (body length and mass) is one of the major axes of functional space (Carmona et al. 2021). It is included in most, if not all, analyses of functional or ecological distinctiveness (e.g., Leitão et al., 2016; Cooke et al., 2020; Carmona et al., 2021; Burner et al., 2022). Considering body size in distinctiveness measurement places an emphasis on large species because they are less numerous than small species and, in vertebrate groups, are often associated with an increased risk of extinction (Toussaint et al., 2021). In scientific studies, threatened large mammals, however, already receive more attention than nonthreatened species, which themselves receive more attention than threatened small mammals (Trimble and van Aarde, 2011). At least in this group, considering body mass when measuring functional distinctiveness could thus tend to highlight species that are already considered by conservation policies. In contrast, we showed in our case study that large species are not the sole species identified as the most functionally distinctive and threatened by our framework. Small-size (e.g., the large rock rat, *C. elvira*, and the kondana rat, *M. kondana*) and medium-size (e.g., the pangolins and the Indian wild dog *C. alpinus*) species were also considered top conservation priorities, even when only body mass was considered to characterize the species.

Trait selection depends on the taxonomic group, spatial scale and habitat considered. Previous studies on functional distinctiveness often included several traits across four categories: morphology, trophic ecology (feeding), reproduction, behavior and habitat preferences (e.g., Cooke et al., 2020; Soares et al., 2021; Trindade-Santos et al., 2022). In our case study, we focused first on body mass and then included available information on trophic ecology, behavior (activity cycle) and habitat preferences, for which we did not have to impute data. Studies are indeed generally limited by trait availability (e.g., Carmona et al., 2021), leading to partial views of the functional distinctiveness of a species. Large-scale studies often have to impute trait data, considering correlations with other available traits or the trait values of related species (e.g., Cooke et al., 2020).

Apart from body mass, we did not include morphometric traits. Indeed, it is still unclear whether species with odd morphologies have rare ecological roles (Redding et al. 2010). This might depend on the group and on the morphometrical traits considered. For example, morphological traits can be used to inform the mobility mode of a species (e.g., Trindade-Santos et al., 2022). They are also often used to precisely determine the diet of the species. It is well known, for instance, that data on beak shape are useful to characterize the functional role of birds, notably in terms of seed dispersal (e.g., Wheelwright, 1985). The shape of fish teeth is also related to the type of consumed items (e.g., Soares et al., 2021, and references therein).

In addition, we focused on effect traits in our case study. In contrast to other studies (e.g., Cooke et al., 2020; Carmona et al., 2021; Toussaint et al., 2021; Burner et al., 2022), we did not include life history traits linked to reproduction. Reproductive traits could be qualified as effect traits by considering, for instance, that they influence food availability (e.g., for prey) (Luck et al., 2012) and the level of pressure on other organisms (e.g., for predators). However, reproductive traits could be more directly linked to the persistence strategies of species and could thus be more directly viewed as response traits. For instance, species with a slow pace of life tend to be threatened with extinction (Toussaint et al., 2021). Additionally, reproduction was partially included in our model of extinction probabilities via the consideration of generation length.

In addition to functional traits, our approach considered each species’ extinction probability. It is thus sensitive to the way species extinction probabilities are quantified and is also limited by the availability of data on extinction probabilities. Access to precise estimates of local extinction probabilities relying on models in population genetics and population dynamics are not yet available for such a large set of species as all mammals found in a rich region. We thus used the IUCN red list to obtain a global index of species extinction risk. Although regional definitions of the IUCN categories of extinction risk have started to be published (IUCN, 2022), they are limited to a few geographic areas (e.g., several African regions) and a few taxonomic groups of interest for conservation, with the European region having the highest number of regionally evaluated species. However, global and regional assessments may be different. For example, the lion *P. leo* is evaluated globally as vulnerable (IUCN, 2022) but endangered in India (*P. leo spp. persica*, IUCN 2022). Therefore, its extinction probability may have been underestimated in our study. However, we could not find such regional assessments for all species in our case study.

In addition, there are currently no identified consensus models of species extinction probability (e.g., Mooers et al., 2008). The model (Andermann et al., 2021) we selected conforms to the IUCN definition of the extinction probabilities of endangered and critically endangered species (IUCN, 2022) that are interpreted with regard to the generation length: an endangered species may have a higher extinction risk than a critically endangered species if its generation length is far shorter. Species with long generation lengths tend to be large species. Although several large-bodied species are classified as critically endangered in our data set and within mammals more globally (Cardillo et al., 2005), their extinction probabilities within a short time frame (<100 years) are thus not the highest.

### 5.2. Choosing a single viewpoint on functional distinctiveness influences conservation priorities

The second step when studying functionally distinctive and threatened species is to select a relevant formula to quantify to what degree a species is functionally distinctive and threatened. This quantitative measurement will then serve to rank species from the most to the least functionally distinctive and threatened. Choosing a formula means choosing precisely what we want to conserve in priority. For example, according to Kosman et al. (2019), “a species might be, on average, quite distant from most species but functionally similar to one other species. The conservation priority for such a species might be lower than that of a species with the same average distance to other species, but that is also consistently different from all other species (i.e., a species without a close neighbour in functional space)”.

According to Kosman et al. (2019), distinctive species are worth increased consideration in conservation practices even if they are currently widespread and not threatened. The measurement of distinctiveness is thus key to detecting these species and either planning or reinforcing restoration and conservation actions for threatened distinctive species or preventive surveys and actions for least concern or near threatened distinctive species. Looking at various values of *α*, our framework allows navigating among different definitions of species’ distinctiveness, from the minimum to the maximum functional dissimilarity to another species, while accounting (index ^*α*^*f*) or not (^*α*^*Deq*\*) for species’ extinction probabilities. High values of *α* highlight species at the edges of the trait space, with extreme trait values (e.g., for body mass in our case study, the pygmy white-toothed shrew *S. etruscus* weighing less than 3 grams and the elephant *E. maximus* weighing more than 3 tons). Protecting them could help to maintain a broad range of ecological strategies (Cooke et al., 2020). Some of these species may be functionally close. A species can thus be considered distinctive for high but not for low values of *α* (e.g., the shrew *Suncus stoliczkanus* and the mouse *Mus terricolor*, in our case study with body mass close to 9 grams). “Functional outliers” (Violle et al., 2017) could thus be defined as the species that are distinctive regardless of *α*, as they are at the border of the “functional space” and far from all others. As expected in our case study, our framework identified the elephant and rhino as functional outliers when considering body mass only. It identified the vulnerable clouded leopard *N. nebulosa* and the endangered Eld’s deer (Sangai) *R. eldii* as functional outliers when a range of traits was considered.

Similar to others (e.g., Carmona et al., 2021), we connected via the indices ^*α*^*f*′ and ^*α*^*f* ′′ the distribution of functional distinctiveness with that of the species’ own extinction probability to understand the risk of losing unique functional strategies. ^*α*^*f*′ increases with both a species’ functional distinctiveness and its extinction probability, while ^*α*^*f* ′′ increases with both a species’ functional distinctiveness and its survival probability. For two species with similar functional distinctiveness, ^*α*^*f*′ would thus attribute a higher value to the most threatened, while ^*α*^*f* ′′ would attribute a higher value to the least threatened. Our analysis also showed that when a species has a drastically high functional distinctiveness compared to others, it can be classified both in the top ^*α*^*f*′ and in the top ^*α*^*f* ′′ highest values, which reinforces the message that highly functionally distinctive species are worthy of increased consideration in conservation practices even if their extinction probability in the coming years is not the highest. In these indices and in ^*α*^*f*, we also highlighted the importance of including the extinction probabilities of other species when assessing the functional distinctiveness of a target species. This is because a species is all the more distinctive that its lookalikes, in terms of functional traits, are likely to be lost in the near future, which implies that its trait values are threatened with disappearance. As such, the loss of functionally distinctive species would thus mean a large effective or potential loss in functional diversity.

### 5.3. Potential consequences of the loss of functionally distinctive species

Once the degree to which a species is functionally distinctive and threatened has been measured, such a measurement can be integrated as a criterion in conservation planning because functionally distinctive species are assumed to have irreplaceable roles in their ecosystem. By definition, the loss of unique species that are functionally different from all other species would lead to functional clustering and thus low functional diversity. Such a correlation between threat and distinctiveness was observed, for example, by Cooke et al. (2020) in birds and mammals. At a global level, Carmona et al. (2021) observed that threatened mammals tend to be close to the boundaries of trait distributions. Assuming that rare species are likely the first locally threatened species in the case of human-driven disturbance, Leitão et al. (2016) for fish and Burner et al. (2022) for beetles also observed a link between functional distinctiveness and extinction risk.

Enhanced conservation of distinctive species is thus needed to avoid the decrease in functional diversity and the deterioration of ecosystem function (Brodie et al., 2021). There is indeed a growing body of evidence that threatened and distinctive species are keystone species, i.e., are critical for the integrity of their ecosystem. Large-bodied mammal species, for example, are ecologically distinctive (Cooke et al., 2020) and keystone species for ecological processes (Toussaint et al., 2021). Some have unique roles as high-level predators, while others are large herbivores (Cooke et al., 2020). In our data set, Asian elephants, for instance, can structure ecosystems by altering vegetation structure and composition (Cooke et al., 2020); large felids (such as the Asiatic population of lion *P. leo* and the tiger *P. tigris*) are notable high-level predators. However, large-bodied mammals with high longevity, late sexual maturity, and long weaning and gestation periods also tend to be among the most threatened (Carmona et al., 2021). In mammals, as in other vertebrate groups, the expected extinction of large species with a slow pace of life could thus particularly lead to potential loss of functional diversity (Toussaint et al., 2021).

Among smaller species, it is striking that, even when considering a single trait (body mass) and a region that is at the border of its distribution range, the Chinese pangolin (*P. pentadactyla*) is classified among the four top distinctive and threatened species of Indian dry forests, followed by the Indian pangolin (*P. crassicaudata*). When also considering a range of functional traits, both species still ranked among the few most distinctive and threatened. Considering a wider range of traits, these species would likely appear even more distinctive, as pangolins notably have unique anatomical and physiological traits (Heighton and Gaubert, 2021). Regarding their roles in their ecosystem, pangolins are prey for large mammals, including tigers and Asiatic lions; they are also predators of insects, contributing to insect (including pest) regulation, and they improve soil quality by excavating burrows (Chao et al., 2020).

The functional diversity of mammals is thus threatened by the current patterns of extinction risks in this group. This global decline will involve regional decreases in functional diversity through the loss of functionally distinctive species, a decrease that could, in the long term, lead to cascading extinctions in fragile ecosystems (Carmona et al., 2021).

### 5.4. Fundability of conservation actions for functionally distinctive and threatened species

Conservation actions exist for most species that we identified as distinctive and threatened in the Indian dry forests. They significantly contributed to the preservation of these species but were not sufficient to avoid drastic declines in their populations in the last few decades. The Indian rhinoceros (*R. unicornis*) has experienced a drastic loss of its distribution range (99.7% decrease from 1970 to 2020), and most remaining individuals are now restricted to two protected areas, one in India and the other in Nepal (Pacifici et al., 2020). Protected areas, such as national parks with antipoaching measures and regulation of prey availability (Pacifici et al., 2020), indeed benefited large-bodied threatened species. For example, tigers, which have experienced a large increase in population size in recent decades within Rajiv Gandhi National Park in India (Karanth et al., 2006), have seen their distribution range drastically reduced outside protected areas (Pacifici et al., 2020). For some species (such as the Namdapha flying squirrel *B. biswasi*), conservation actions mainly consist of occurrence in a protected area (IUCN, 2022). Protected areas may be the last chance for these species. However, such areas have restricted ranges (with animal persecution still occurring via illegal trade, especially on the borders) and are often disconnected and isolated (absence of corridors).

Other species (e.g., the large rock rat *C. elvira* and the kondana rat *M. kondana*) have no conservation actions listed by IUCN (2022). For example, *C. elvira* is considered by the Alliance for Zero Extinction (https://zeroextinction.org/; accessed on April 20, 2022) to justify and promote the protection of natural areas of Kurumbapatti village in the Eastern Ghats of India, a small refuge where the remaining declining populations of *C. elvira* live. Our case study thus illustrates that enhanced conservation actions are still necessary for threatened functionally distinctive species (regionally and globally) to avoid extinction in the coming decades.

Conservation initiatives that consider distinctiveness as an important criterion are still rare. Launched in 2007 (Isaac et al., 2007), the “edge of existence” (EDGE) initiative nevertheless collects one-time and monthly donations by the public to support actions for the preservation of evolutionarily distinctive and threatened species in many countries around the world. Similar to what is done by the EDGE initiative, other initiatives could identify species that are threatened and functionally distinctive but have not or have rarely attracted the attention of conservation initiatives. Indeed, discrepancies between functional and evolutionary distinctiveness have been observed (e.g., Cooke et al., 2020). In addition, in contrast to the analysis of evolutionary distinctiveness, which is currently global, the study of functional distinctiveness could be carried out, as in our case study, more locally within countries and within habitat types, to more directly characterize the impact the loss of a species could have on ecosystem functioning. For example, none of the functionally distinctive squirrels (*B. biswasi*) and rats (*C. elvira* and *M. kondana*) that lack conservation actions in Indian dry forests were considered by the EDGE initiative. Their conservation would be worth careful study in light of cultural, economic (source of food for local population for squirrels) and sanitary (for rats) aspects. As conservation actions currently mostly rely on individual donations, the feasibility of such an approach would depend on the motivation of the general public to preserve threatened functionally-distinctive species. This motivation will have to be evaluated in a wide range of taxonomic groups.

Large endangered species are indeed typically used as flagships in conservation marketing campaigns (e.g., Macdonald et al., 2015; and references therein). Cooke et al. (2020) found that ecologically distinctive mammal species tend to be charismatic species. Additionally, the most charismatic species for the public tend to be large-size species with high (threatened) IUCN scores (Macdonald et al., 2015), especially mammals (Albert et al., 2018), and many of them have populations in India: the tiger (*P. tigris*), the lion (*P. leo*), the elephant (including *E*. maximus), the leopard (*P. pardus*), the rhino (including *R. unicornis*) and the gray wolf (*Canis lupus*).

Felids and primates are highly favored species by the general public (including Indian people) for funding conservation actions, with the tiger being the most favored by a wide margin (Macdonald et al., 2015). Preserving umbrella species with large home ranges and habitat requirements can have cobenefits on the conservation of cooccurring species and their ecosystems. This is the case for most charismatic species (Courchamp et al., 2018) and especially for the tiger (Vasudeva et al., 2022). Our approach, applied to a range of functional traits, highlighted 3 felids (the tiger *P. tigris* and the leopards *P. pardus* and *N. nebulosa*) and 1 primate (the gibbon *H. hoolock*) among the most functionally distinctive and threatened species in Indian dry forests. The fact that a species is charismatic can motivate the general public to fund conservation actions (Albert et al., 2018; Courchamp et al., 2018). However, despite conservation actions, many charismatic species are still declining (Courchamp et al., 2018; IUCN, 2022).

Even within charismatic taxonomic groups, such as mammals, the feasibility of conservation campaigns that target noncharismatic species still has to be determined. For example, pangolins have been highlighted in our data set as threatened functional outliers in Indian dry forests when accounting for their and other species’ extinction probabilities. Pangolin species are well known by conservation actors to be poached, being the species most heavily trafficked to extinction among wild mammals. Despite this, they only recently attracted the interest of the media because the pangolin has been, without concrete evidence, claimed to be an intermediate host of SARS-CoV-2 at the origin of the COVID-19 pandemic (Heighton and Gaubert, 2021), an increased interest that could, however, lead to a negative public perception of the interest in preserving pangolins (Choo et al., 2022).

### 5.5 Conclusion

We provide herein a unified parametric framework for the measurement of functional distinctiveness. In this framework, functional distinctiveness is simply measured as a weighted mean of functional differences with other species. The framework is said to be parametric because the weights in the weighted mean depend on a parameter that allows navigating through different definitions of functional distinctiveness from the minimum to the maximum functional difference from another species. This framework could avoid unconscious bias in this context due to the selection of a single nonparametric formula to measure distinctiveness. Applying this framework to a variety of taxa, habitats and realms will likely reveal a large diversity of noncharismatic, less well-known threatened species with distinct functional traits and ecological strategies that have key roles in their ecosystems. There is thus a need to study the feasibility of such an approach on understudied species and taxonomic groups for which functional traits may be available at regional levels rather than at a global level. We hope our framework, which brings together different viewpoints on how functional distinctiveness must be defined, will help communicate this concept as one of the criteria that should be considered when prioritizing species within countries for conservation.

## Acknowledgements

We thank all experts who contributed to the IUCN Red List. We also thank all authors of silhouette images of animals that have offered their work to PhyloPic (http://phylopic.org) for public domain dedication.

## Author contributions

**Sandrine Pavoine:** Conceptualization, Methodology, Software, Formal analysis, Writing-Original draft preparation, Visualization. **Carlo Ricotta**: Writing-Reviewing and Editing.

## The following are the supplementary data related to this article

Appendix A. Mathematical proofs.

Appendix B. Supplementary analyses done on the mammals of Indian dry forests.

Appendix C. Dataset.

Appendix D. R function.

Appendix E. R script.

