## Appendix A for "Identifying functionally distinctive and threatened species"

### Appendix A. Mathematical proofs

Consider, the same notations as in the main text. In Pavoine and Ricotta (2021) we introduced index ${}^{\alpha}Deq_j^*$ as a parametric index of functional distinctiveness as follows:

$$ {}^{\alpha}Deq_j^* = \frac{\sum_{c=1}^{N-1} u_{c|j}(N^{\alpha-1} - c^{\alpha-1})}{N^{\alpha-1}-1} \quad \text{(A.1)} $$

where $u_{c|j}$ amounts to how much more dissimilar *c* is to *j* than species *c* - 1 is: $u_{c|j} = d_{c,j} - d_{c-1,j}$, with $d_{0,j} = d_{jj} = 0$. We considered ${}^{1}Deq_j^*$ equal to the limit of ${}^{\alpha}Deq_j^*$ when $\alpha$ tends to 1, which yields: ${}^{1}Deq_j^* = \sum_{c=1}^{N-1} u_{c|j}(1 - \log_N(c))$.

#### Proof that Eq. A.1 above and Eq. 1 of the main text are equivalent

We start with Eq. A.1, i.e. ${}^{\alpha}Deq_j^* = \frac{\sum_{c=1}^{N-1} u_{c|j}(N^{\alpha-1} - c^{\alpha-1})}{N^{\alpha-1}-1}$

where *c* is the *c*$^{th}$ species most functionally similar to *j*, $d_{c,j}$ its dissimilarity with species *j* (posing $d_{0,j} = d_{jj} = 0$) and $u_{c|j} = d_{c,j} - d_{c-1,j}$ (Pavoine and Ricotta, 2021).

$$ \sum_{c=1}^{N-1} u_{c|j}(N^{\alpha-1} - c^{\alpha-1}) = \sum_{c=1}^{N-1} u_{c|j} \sum_{m=c}^{N-1}((m+1)^{\alpha-1} - m^{\alpha-1}) $$

$$ = \sum_{m=1}^{N-1}((m+1)^{\alpha-1} - m^{\alpha-1}) \sum_{c=1}^{m} u_{c|j} $$

$$ = \sum_{m=1}^{N-1}((m+1)^{\alpha-1} - m^{\alpha-1})\, d_{m,j} $$

□

**Proof that $\sum_{c=1}^{N-1} \pi_c = 1$ (Eq. 2 of the main text):**

$$\pi_c = \frac{(c+1)^{\alpha-1} - (c)^{\alpha-1}}{N^{\alpha-1} - 1}$$

$$\sum_{c=1}^{N-1} \pi_c = \sum_{c=1}^{N-1} \frac{(c+1)^{\alpha-1} - (c)^{\alpha-1}}{N^{\alpha-1} - 1}$$

$$= \frac{\sum_{c=1}^{N-1}(c+1)^{\alpha-1} - \sum_{c=1}^{N-1}(c)^{\alpha-1}}{N^{\alpha-1} - 1} = \frac{\sum_{c=2}^{N}(c)^{\alpha-1} - \sum_{c=1}^{N-1}(c)^{\alpha-1}}{N^{\alpha-1} - 1}$$

$$= \frac{(N)^{\alpha-1} + \sum_{c=2}^{N-1}(c)^{\alpha-1} - \sum_{c=2}^{N-1}(c)^{\alpha-1} - (1)^{\alpha-1}}{N^{\alpha-1} - 1} = 1$$

□
