## Appendix B for "Identifying functionally distinctive and threatened species"

**Appendix B. Supplementary analyses done on the mammals of Indian dry forests**

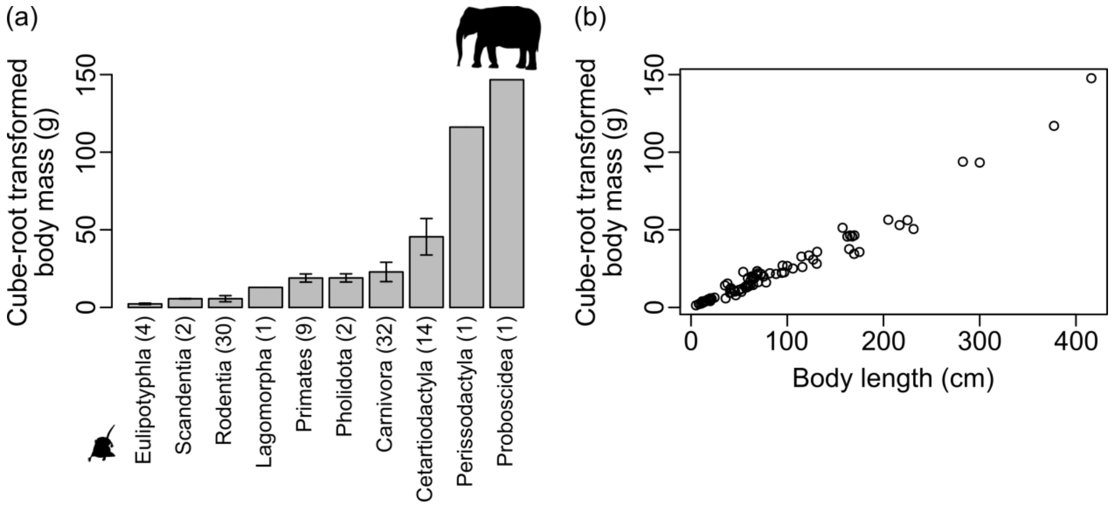


**Fig. B.1.** Information on mammal body mass in the dry forests of India. (a) Mean (grey bars) and standard deviation (segments) of body masses for each mammal order. (b) Link between body mass and body length.

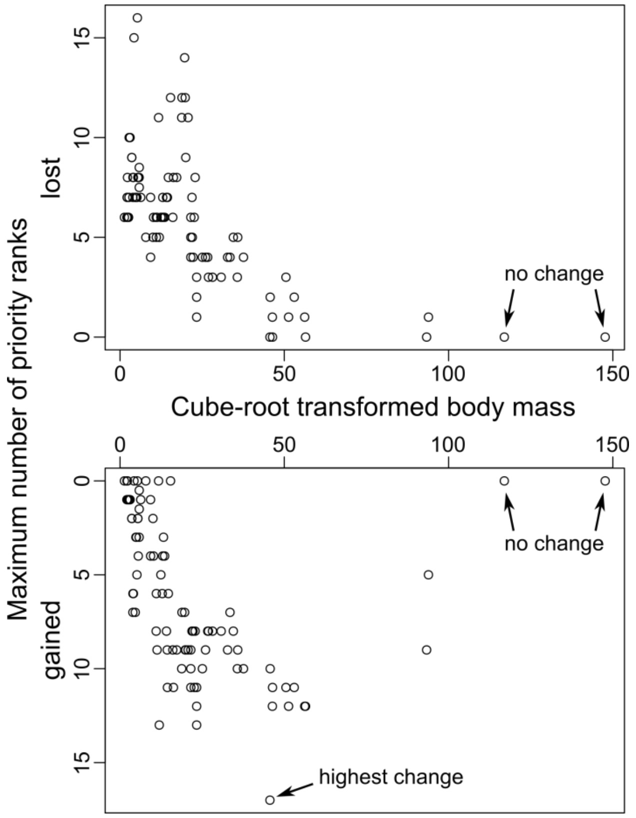


**Fig. B.2.** Maximum number of priority ranks lost or gained by a species when classified according to the ${}^{\alpha}Deq^{*}$ index compared to the ${}^{\alpha}f$ index from the most (1st priority rank) to the least (95th priority rank) distinctive species (considering body mass only). Here species are organized on the x axis according to their body mass.
